# hicap: *in silico* serotyping of the *Haemophilus influenzae* capsule locus

**DOI:** 10.1101/543454

**Authors:** Stephen C. Watts, Kathryn E. Holt

## Abstract

*Haemophilus influenzae* exclusively colonises the human nasopharynx and can cause a variety of respiratory infections as well as invasive diseases including meningitis and sepsis. A key virulence determinant of *H. influenzae* is the polysaccharide capsule of which six serotypes are known, each encoded by a distinct variation of the capsule biosynthesis locus (*cap*-a to *cap*-f). *H. influenzae* type b (Hib) was historically responsible for the majority of invasive *H. influenzae* disease and prevalence has been markedly reduced in countries that have implemented vaccination programs targeting this serotype. In the postvaccine era, non-typeable *H. influenzae* emerged as the most dominant group causing disease but in recent years a resurgence of encapsulated *H. influenzae* strains has also been observed, most notably serotype a. Given the increasing incidence of encapsulated strains and the high frequency of Hib in countries without vaccination programs, there is growing interest in genomic epidemiology of *H. influenzae*. Here we present hicap, a software tool for rapid *in silico* serotype prediction from *H. influenzae* genome sequences. hicap is written using Python3 and is freely available at https://github.com/scwatts/hicap under the GPL3 license. To demonstrate the utility of hicap, we used it to investigate the *cap* locus diversity and distribution in 691 high-quality *H. influenzae* genomes from GenBank. These analyses identified *cap* loci in 95 genomes and confirmed the general association of each serotype with a unique clonal lineage and also identified occasional recombination between lineages giving rise to hybrid *cap* loci (2% of encapsulated strains).

## Introduction

*Haemophilus influenzae* is a pleomorphic Gram-negative bacterium that is exclusive to humans, typically colonising the upper respiratory tract and occasionally causing disease. It was the first free living organism to be completely sequenced and served as a stepping stone towards DNA sequencing technology development in preparation for the Human Genome Project [1]. *H. influenzae* is often classified on the basis of the production and antigenicity of polysaccharide capsule. Strains that produce capsule are divided into six serotypes (Hia to Hif), and non-encapsulated strains are designated non-typeable *H. influenzae* (NTHi) [2].

Biosynthesis of the polysaccharide capsule is controlled by the *cap* loci (*cap*-a to *cap*-f), which each include three contiguous but functionally distinct regions (I, II, and III) (Figure 1). Regions I and III are common to all six *cap* loci, and are associated with cellular transport (*bex* operon, region I) and post-translational processing (*hcs* operon, region III) [3, 4]. Region II encodes several genes involved in polysaccharide biosynthesis that are specific to each serotype (Figure 1) [5–9]. The *cap* locus is regularly subject to duplication, deletion, and interruption [10, 11]. For example the *cap*-b locus is often duplicated, creating two tandem copies of the locus flanked by IS*1016* and regularly coincides with a 1.2 kbp deletion of the terminal *bexA*-IS*1016* copy [9]. The arrangement and copy number of *cap* locus genes also has clinical relevance as certain structural variants are associated with increased levels of virulence [12].

**Fig 1.**
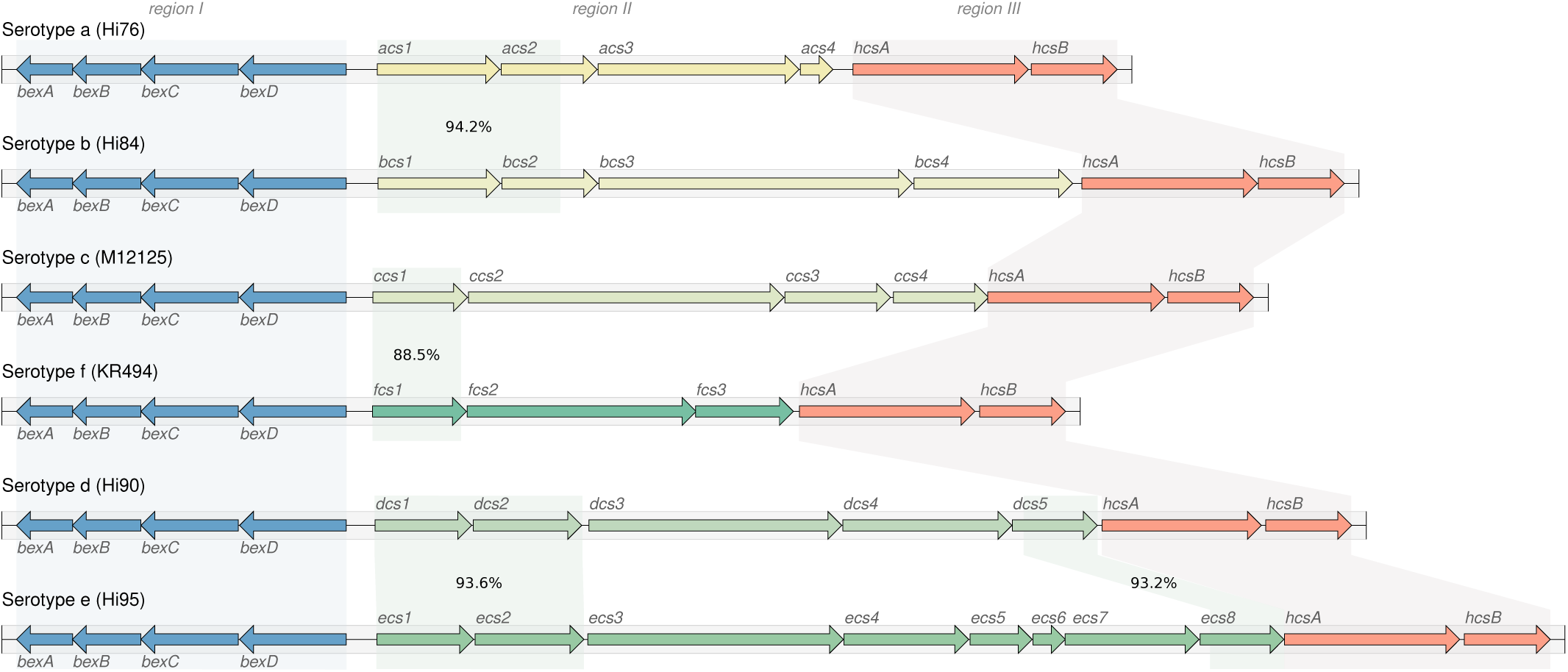
Schematic representation of the six known *H. influenzae cap* loci. Capsule nucleotide sequences and annotations were collected from genome assemblies representing each of the six serotypes. Shading indicates homologous regions between reference loci as determined by BLAST identity values shown in Region II. Region I and III are homologous across the entire sequence for all loci with a nucleotide identity of ≥ 87% and ≥ 90% respectively.

*H. influenzae* is capable of causing a variety of respiratory infections and invasive diseases. Prior to the introduction of capsular conjugate vaccines against Hib in the 1980s, this serotype was responsible for almost all *H. influenzae* related morbidity and mortality [13]. In the period subsequent to wide-spread adoption of childhood Hib vaccination programs, the incidence of Hib-related disease reduced markedly [14]. However following implementation of Hib vaccination programs, disease caused by NTHi has been increasing globally. Disease prevalence of other encapsulated strains is also increasing at an alarming rate and Hia disease burden now exceeds that of Hib during the pre-Hib vaccination era in some regions and populations [15]. A recent report of particular interest found that Hia constituted 50% of all *H. influenzae* cases between 2010 and 2015 in Northwestern Ontario, Canada [16]. Importantly Hib also remains an issue in countries that have not implemented a vaccination program [17].

Public health and clinical laboratories are now beginning to incorporate whole genome sequencing (WGS) technologies into diagnostic, outbreak, and surveillance programs [18, 19]. The departure from molecular based diagnostics has been largely driven by considerably higher resolution and accuracy afforded by WGS [20]. Currently there are no dedicated tools for *H. influenzae* serotype prediction that seek to leverage WGS data for *cap* loci detection. The need for such a tool continues to grow with the resurgence of encapsulated *H. influenzae* and the increasingly routine use of WGS in the public health setting.

Here we describe hicap, a software tool specifically designed for rapid *in silico* serotype prediction from *H. influenzae* WGS data. hicap is an open source Python3 package and is freely available at https://github.com/scwatts/hicap under a GPLv3 license. We further apply hicap to identify and extract *cap* loci from all *H. influenzae* genomes currently available in GenBank, and explore diversity and distribution of these loci in the *H. influenzae* population.

## Methods

### hicap implementation and validation

hicap uses a reference database to identify genes expected in the six *cap* loci (*cap*-a to *cap*-f). To this end, a curated nucleotide sequence database of *cap* locus genes was constructed by extracting the protein-coding sequences annotated from *cap* loci in publicly available sequences of well defined *H. influenzae* serotypes (Table 1). Using this database, the process adopted by hicap to perform serotype prediction from WGS assemblies is described in Figure 2.

**Table 1.** *cap* locus sequences used to create the hicap reference database.

| Gene/ Region | Strain | Accession | Reference |
| --- | --- | --- | --- |
| *bexA* | KR494 | GCA_000465255.1 | [42] |
| *bexB* | KR494 | GCA_000465255.1 | [42] |
| *bexC* | KR494 | GCA_000465255.1 | [42] |
| *bexD* | KR494 | GCA_000465255.1 | [42] |
| *cap-a* | Hi76 | ERX1834399 | [43] |
| *cap-b* | NCTC 8468 | ERX704106 | [44] |
| *cap-c* | Hi85 | ERX1834408 | [43] |
| *cap-d* | ATCC 9008 (*cap* locus) | HQ424464.1 | *Haemophilus influenzae* ATCC 9008™ |
| *cap-e* | hi467 | GCA_001975845.1 | [45] |
| *cap-f* | KR494 | GCA_000465255.1 | [42] |
| *hcsA* | KR494 | GCA_000465255.1 | [42] |
| *hcsB* | KR494 | GCA_000465255.1 | [42] |
| IS1016 | Hae18 | X59756.1 | [46] |

**Fig 2.**
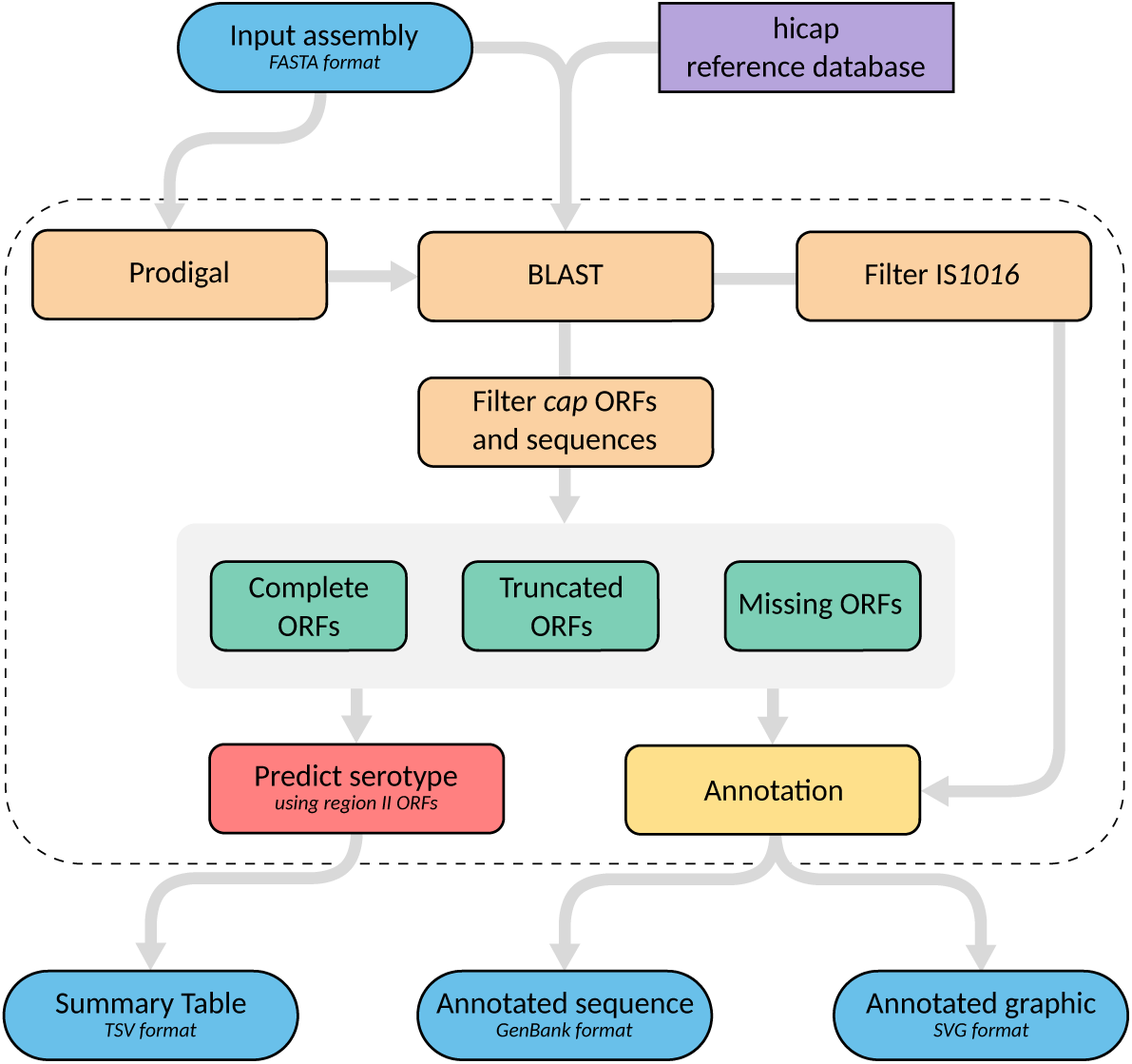
Summary of the hicap serotype prediction method. hicap takes an assembled genome in FASTA format as input and detects all open reading frames (ORFs) using Prodigal. Constituent *cap* genes and IS*1016* copies are identified by performing alignments of either ORF sequence or input assembly sequence against the reference database using BLAST+. The identified *cap* genes and IS*1016* alignments are then used to inform structural composition of the locus. Serotype is predicted using the gene complement information of region II.

Firstly all open reading frames (ORFs) are identified in the query assembly using Prodigal [21]. Then each ORF nucleotide sequence is queried against the hicap reference database using BLAST+ [22]. The resulting alignments are filtered on the basis of subject coverage and nucleotide identity. The default parameters to designate an ORF as a complete match to a *cap* locus gene are subject coverage ≥ 80% and nucleotide identity ≥ 70%. Often *cap* genes that are expected to be present lack a complete match to any ORF annotated by Prodigal. This typically occurs when an ORF in the Prodigal annotation has been truncated due to missense mutations, mobile elements, or incomplete assembly. hicap infers the number of genes missing from the predicted *cap* locus by examining the count of complete ORFs and comparing this to the expected count for the complete form of that locus.

Generally hicap will attempt to find at least one copy of each gene expected in the *cap* locus. In the case that there are missing genes, hicap searches the remaining ORF-database alignments for the expected gene fragment(s) using more relaxed filtering (defaults for this are alignment length ≥ 60 bp and nucleotide identity ≥ 80%). Failing this, hicap employs BLAST+ to identify regions of the input assembly that are homologous to missing genes proximal to the predicted *cap* locus (filtering alignments with a bitscore ≤ 200). An ORF or sequence is designated as truncated if it is identified by either of these adjusted filters but not meeting the criteria for a complete match. Additionally hicap searches for IS*1016* in the *cap* locus and nearby regions by aligning the reference IS*1016* sequence with the input assembly using BLAST+.

The resulting set of alignments and ORFs are used to predict serotype and various locus characteristics. Specifically, hicap predicts serotype by considering all complete and truncated alignments of region II genes. The predicted serotype is defined as the serotype observed to have the most complete set of region II genes. Where an ORF has multiple alignments to the hicap database, a single best alignment is selected on the basis of E-value with ties broken by bitscore. ORFs identified as belonging to the *cap* locus and surrounding region are summarised in a tab-delimited report file, and the annotated *cap* locus sequence is output in GenBank format. A visualisation of the locus annotation is also created using the graphics module in Biopython [23] and output in SVG format (examples in Figure 3).

**Fig 3.**
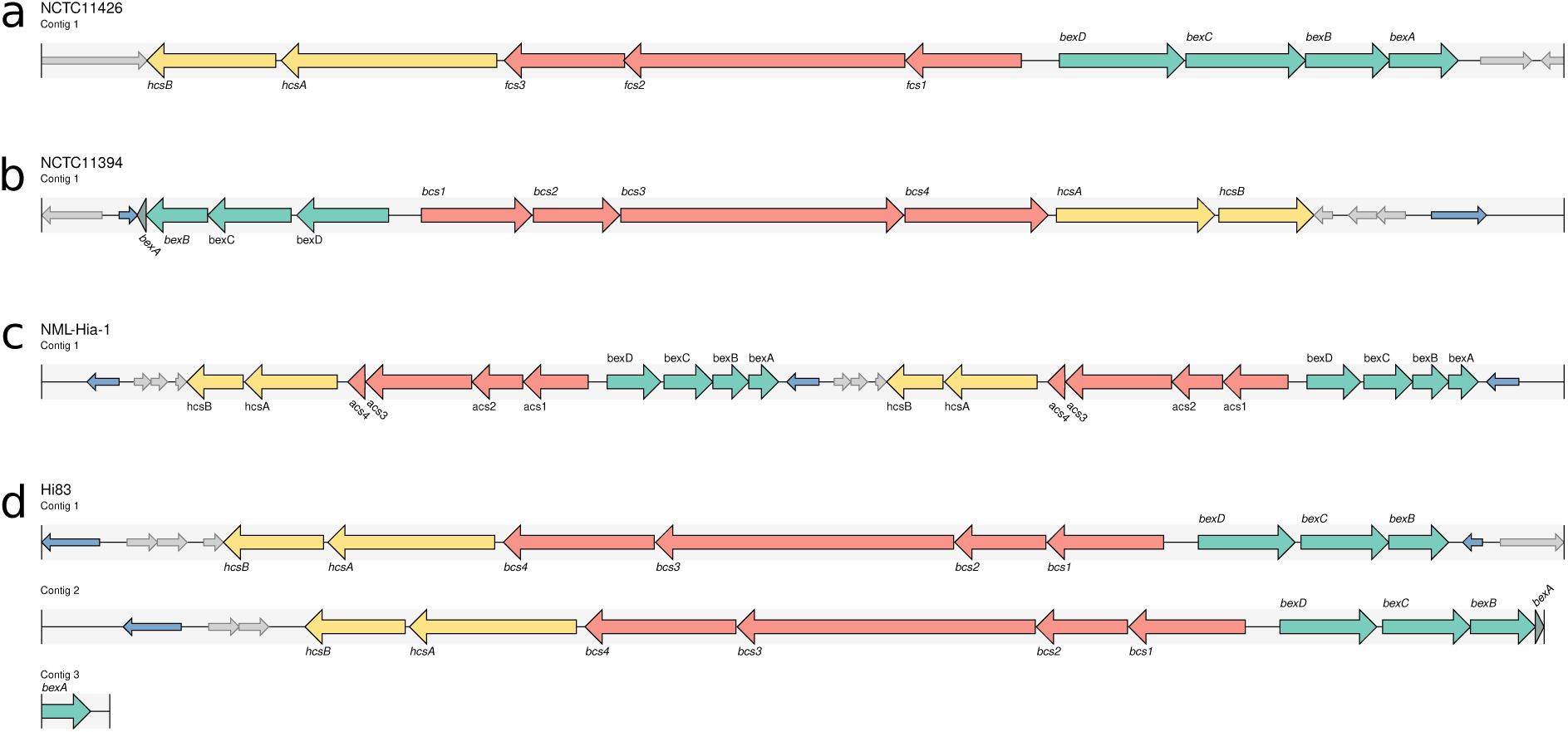
Example hicap visualisation for selected genomes. *cap* locus genes are annotated as large arrows with direction representing strand. Genes of the *cap* locus are coloured to indicate region (region I: green; region II: red; region III: yellow). A truncated *cap* gene is given a darker colour of the respective region. Copies of IS*1016* are denoted as small blue arrows and open reading frames that do not generally belong to the *cap* locus are show as small grey arrows. (a) The complete and contiguous annotation of NCTC 11426 *cap*-f locus. (b) The NCTC 11394 *cap*-b locus that contains a truncated *bexA* gene and two copies of IS*1016*. (c) A duplication of the *cap*-a locus is observed in the assembly of NML-Hia-1. (d) The *cap*-b locus of Hi83 is also duplicated but present across multiple contigs in the input assembly, as represented by multiple tracks.

To test the ability of hicap to predict serotypes, we reviewed the literature and identified publicly available WGS data for *H. influenzae* isolates with known serotype (Table 2). The genome assemblies were downloaded and analysed using hicap run with default parameters. For 26 isolates only read data was available, hence *de novo* assemblies were generated using SPAdes v3.12.0 [24] prior to analysis with hicap. All validation was performed using hicap v1.0.0. The full set of assemblies used for testing are available in FigShare (https://doi.org/10.26180/5c352c5110712). When discrepancies were observed between expected serotype and hicap results, the output of nucleotide BLAST+ v2.7.1+ searches of genome assemblies against the hicap database were manually inspected.

**Table 2.**
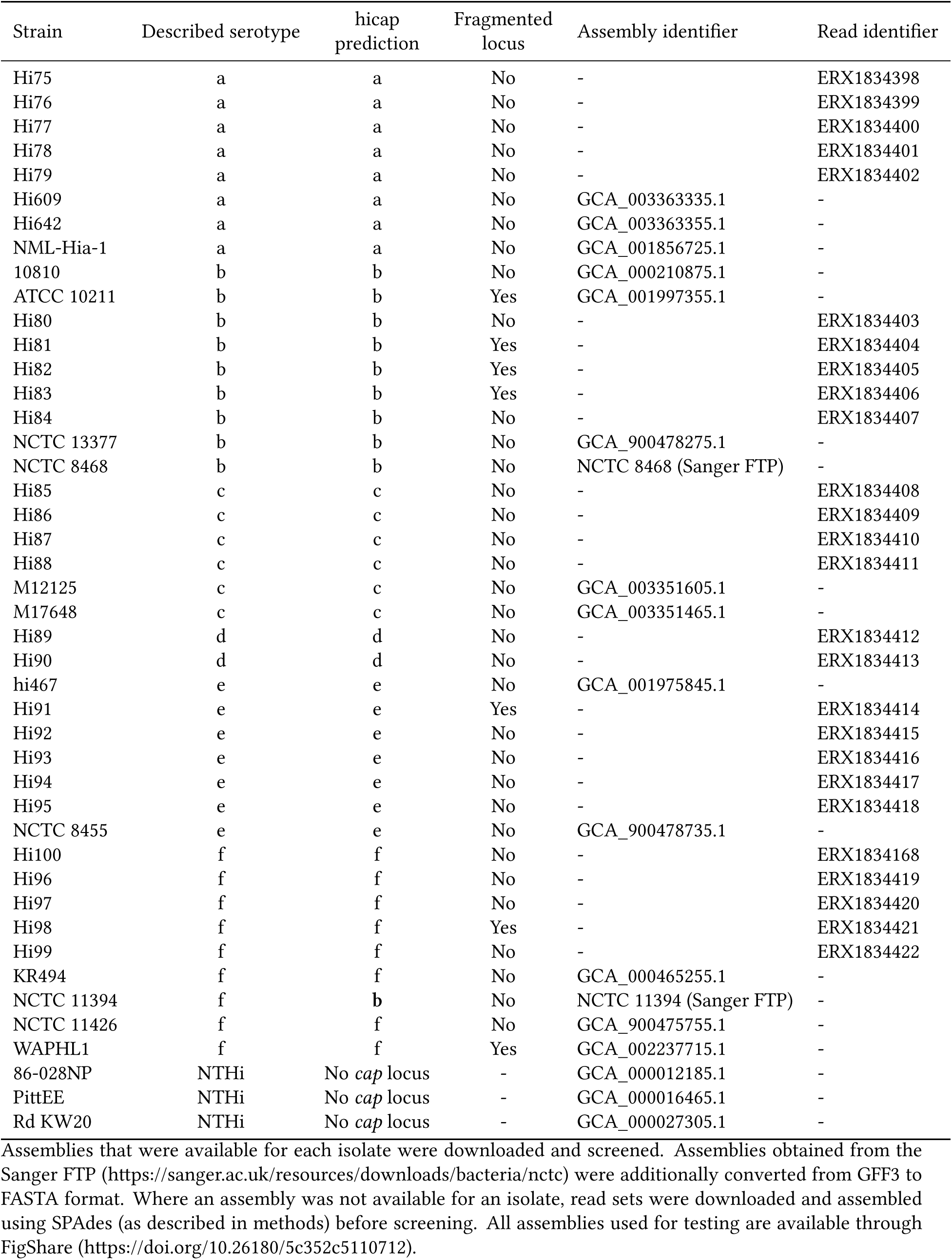
Strains used in the validation of the hicap method with predicted serotype and fragmentation status of the *cap* locus as determined by hicap.

| Strain | Described serotype | hicap prediction | Fragmented locus | Assembly identifier | Read identifier |
| --- | --- | --- | --- | --- | --- |
| Hi75 | a | a | No | - | ERX1834398 |
| Hi76 | a | a | No | - | ERX1834399 |
| Hi77 | a | a | No | - | ERX1834400 |
| Hi78 | a | a | No | - | ERX1834401 |
| Hi79 | a | a | No | - | ERX1834402 |
| Hi609 | a | a | No | GCA_003363335.1 | - |
| Hi642 | a | a | No | GCA_003363355.1 | - |
| NML-Hia-1 | a | a | No | GCA_001856725.1 | - |
| 10810 | b | b | No | GCA_000210875.1 | - |
| ATCC 10211 | b | b | Yes | GCA_001997355.1 | - |
| Hi80 | b | b | No | - | ERX1834403 |
| Hi81 | b | b | Yes | - | ERX1834404 |
| Hi82 | b | b | Yes | - | ERX1834405 |
| Hi83 | b | b | Yes | - | ERX1834406 |
| Hi84 | b | b | No | - | ERX1834407 |
| NCTC 13377 | b | b | No | GCA_900478275.1 | - |
| NCTC 8468 | b | b | No | NCTC 8468 (Sanger FTP) | - |
| Hi85 | c | c | No | - | ERX1834408 |
| Hi86 | c | c | No | - | ERX1834409 |
| Hi87 | c | c | No | - | ERX1834410 |
| Hi88 | c | c | No | - | ERX1834411 |
| M12125 | c | c | No | GCA_003351605.1 | - |
| M17648 | c | c | No | GCA_003351465.1 | - |
| Hi89 | d | d | No | - | ERX1834412 |
| Hi90 | d | d | No | - | ERX1834413 |
| hi467 | e | e | No | GCA_001975845.1 | - |
| Hi91 | e | e | Yes | - | ERX1834414 |
| Hi92 | e | e | No | - | ERX1834415 |
| Hi93 | e | e | No | - | ERX1834416 |
| Hi94 | e | e | No | - | ERX1834417 |
| Hi95 | e | e | No | - | ERX1834418 |
| NCTC 8455 | e | e | No | GCA_900478735.1 | - |
| Hi100 | f | f | No | - | ERX1834168 |
| Hi96 | f | f | No | - | ERX1834419 |
| Hi97 | f | f | No | - | ERX1834420 |
| Hi98 | f | f | Yes | - | ERX1834421 |
| Hi99 | f | f | No | - | ERX1834422 |
| KR494 | f | f | No | GCA_000465255.1 | - |
| NCTC 11394 | f | **b** | No | NCTC 11394 (Sanger FTP) | - |
| NCTC 11426 | f | f | No | GCA_900475755.1 | - |
| WAPHL1 | f | f | Yes | GCA_002237715.1 | - |
| 86-028NP | NTHi | No *cap* locus | - | GCA_000012185.1 | - |
| PittEE | NTHi | No *cap* locus | - | GCA_000016465.1 | - |
| Rd KW20 | NTHi | No *cap* locus | - | GCA_000027305.1 | - |

Assemblies that were available for each isolate were downloaded and screened. Assemblies obtained from the Sanger FTP (https://sanger.ac.uk/resources/downloads/bacteria/nctc) were additionally converted from GFF3 to FASTA format. Where an assembly was not available for an isolate, read sets were downloaded and assembled using SPAdes (as described in methods) before screening. All assemblies used for testing are available through FigShare (https://doi.org/10.26180/5c352c5110712).

### *cap* locus distribution, variation, and recombination

To demonstrate a practical application of hicap, we investigated the distribution of capsular serotypes predicted by hicap amongst all *H. influenzae* genomes available in NCBI GenBank (as of October 8, 2018; n=698, listed in Supplementary File S1). Whole genome assemblies were downloaded via FTP and a phylogeny was constructed using mashtree v0.33 (https://github.com/lskatz/mashtree, [25]). Genomes were excluded from analysis where assembly length was more than four standard deviations from the mean or genomic content was sufficiently dissimilar to *H. influenzae* (n=7). Specifically genomic content was assessed by simulating 50,000 error-free reads using wgsim v0.3.1-r13 (https://github.com/lh3/wgsim), which were taxonomically classified by centrifuge v1.0.4-beta [Kim2016] and samples with ≤ 80% *H. influenzae* reads excluded.

Sequence type (ST) for each assembly was determined via comparison to the multi-locus sequence typing (MLST) database for *H. influenzae* (https://pubmlst.org/hinfluenzae) [26] using mlst v2.15 (https://github.com/tseemann/mlst). Capsular serotypes were inferred using hicap v1.0.0 with the default settings. Nucleotide sequence homology between hybrid *cap* loci was assessed by BLAST+ v2.7.1+ and visualised using genoPlotR v0.8.7 [27] in R v3.4.4 [28]. The mashtree phylogeny was annotated with ST and predicted capsular serotype in R v3.4.4 using ggtree v1.12.7 [29].

To establish the relationship between capsular serotype and allelic variants of genes encoded in regions I and III, we constructed individual gene trees. Nucleotide sequences were extracted for all complete region I and III genes that were detected by hicap during analysis of the *H. influenzae* GenBank dataset. For each individual gene, nucleotide sequences were aligned using MAFFT v7.407 with default settings [30] and phylogenies inferred from the alignment using FastTree v2.1.10 with the general time-reversible substitution model [31]. Nucleotide divergence was calculated using ape v5.2 [32] in R v3.4.4 from gene nucleotide alignments.

## Results and discussion

### hicap validation

To validate hicap as a tool for *in silico* serotyping, we analysed 41 publicly available *H. influenzae* genomes with reported serologically confirmed capsule types including representatives for each of the six serotypes and three NTHi (Table 2). The results show that hicap robustly identifies the *H. influenzae cap* locus even in highly discontiguous assemblies. For each *cap* locus the completeness, presence of truncated genes, duplication, contiguity, and serotype were correctly reported.

Capsule loci were detected by hicap in 41/41 genomes with reported serotypes and 0/3 in serologically determined NTHi genomes (Table 2). The predicted serotype matched the reported serotype in 40/41 cases (98%) (Table 2). We found that hicap yielded accurate predictions even from draft genomes where the *cap* locus was fragmented across multiple contigs (observed in 7 genomes from the validation set). Examples of the *cap* loci identified and visualised using hicap are shown in Figure 3.

The single genome with a discrepancy between predicted and reported serotype, NCTC 11394, is described as Hif in the National Collection of Type Cultures (NCTC) but was confidently assigned Hib by hicap analysis of the completed PacBio genome assembly (Figure 3b). Manual assessment of the *cap* locus in the NCTC 11394 genome assembly additionally confirmed the presence of a complete *cap*-b locus (uninterrupted, in a single contig) and absence of any *cap*-f region II genes with all expected *cap*-b protein-coding genes present at ≥ 95% coverage and ≥ 84% homology to those annotated in the *cap*-b reference sequence (excluding a truncated *bexA* gene). The standard slide agglutination test classically used for serological typing of *H. influenzae* has been shown to lack specificity and has been estimated to yield incorrect results at a rate of 17.5% [33]. It therefore appears likely this discordance is due to inaccuracies in the described serotype rather than misidentification by hicap.

### *cap* locus distribution and variation

To demonstrate the utility of *in silico* serotype prediction with hicap, we used it to investigate all publicly available *H. influenzae* genomes that passed quality filtering criteria (n=691, see Supplementary File S1). hicap identified a complete *cap* locus in 95/691 (13.7%) genomes (8 *cap*-a, 54 *cap*-b, 4 *cap*-c, 1 *cap*-d, 20 *cap*-e, and 8 *cap*-f). All genomes contained either zero or one *cap* locus type, but duplication events were observed in 15/95 (15.8%) *cap*-positive genomes (14 *cap*-b and 1 *cap*-a).

Duplication of the *cap*-b locus has been frequently reported and is associated with enhanced virulence, conferred by an increased ability to produce capsule [12, 34]. This duplication is thought to be driven by copies of IS*1016* flanking the capsule locus in some isolates. A common variant of the duplicated *cap*-b locus involves the deletion of 1.2 kbp in one copy of region I, resulting in the truncation of *bexA* and IS*1016*. We observed this duplication deletion variant in 14/54 (26.0%) predicted Hib genomes. Complete *cap*-b duplication without truncation of *bexA* was not observed. In addition, hicap identified a single isolate (NML-Hia-1) containing a tandem duplication of the *cap*- a locus. Strains identified to be carrying *cap* loci were not assessed for capsule production however several of these strains are known to synthesis capsule (e.g. 10810 and NML-Hia-1).

To examine the distribution of *cap* loci in the *H. influenzae* population, we constructed a whole genome phylogeny (Figure 4) and inferred STs according to the *H. influenzae* MLST scheme (Table 3). We observed a high degree of exclusivity for STs in regard to predicted capsular serotypes, with each ST containing zero or one *cap* locus serotype.

**Table 3.**
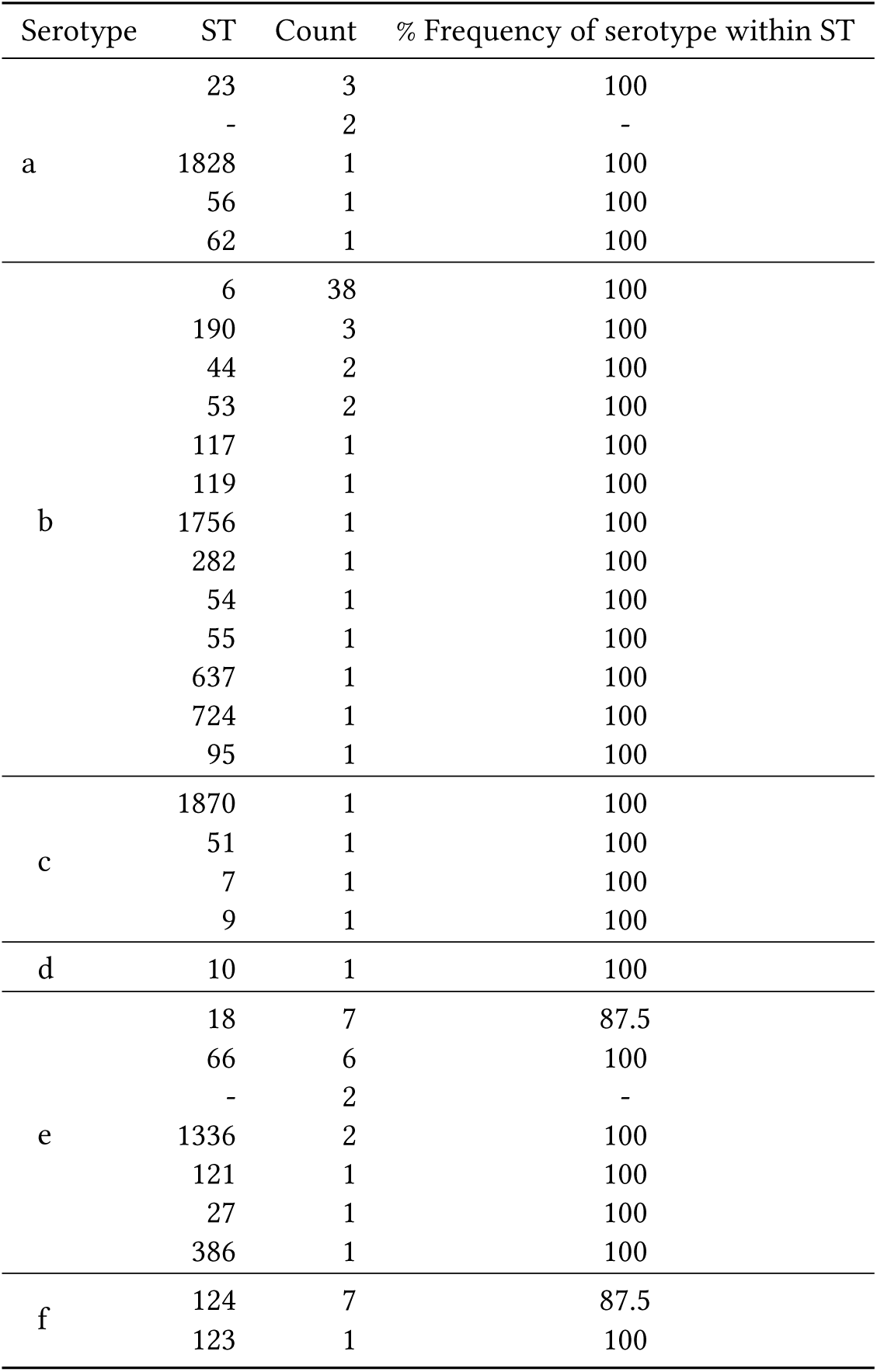
Sequence types associated with each serotype in the GenBank dataset.

**Fig 4.**
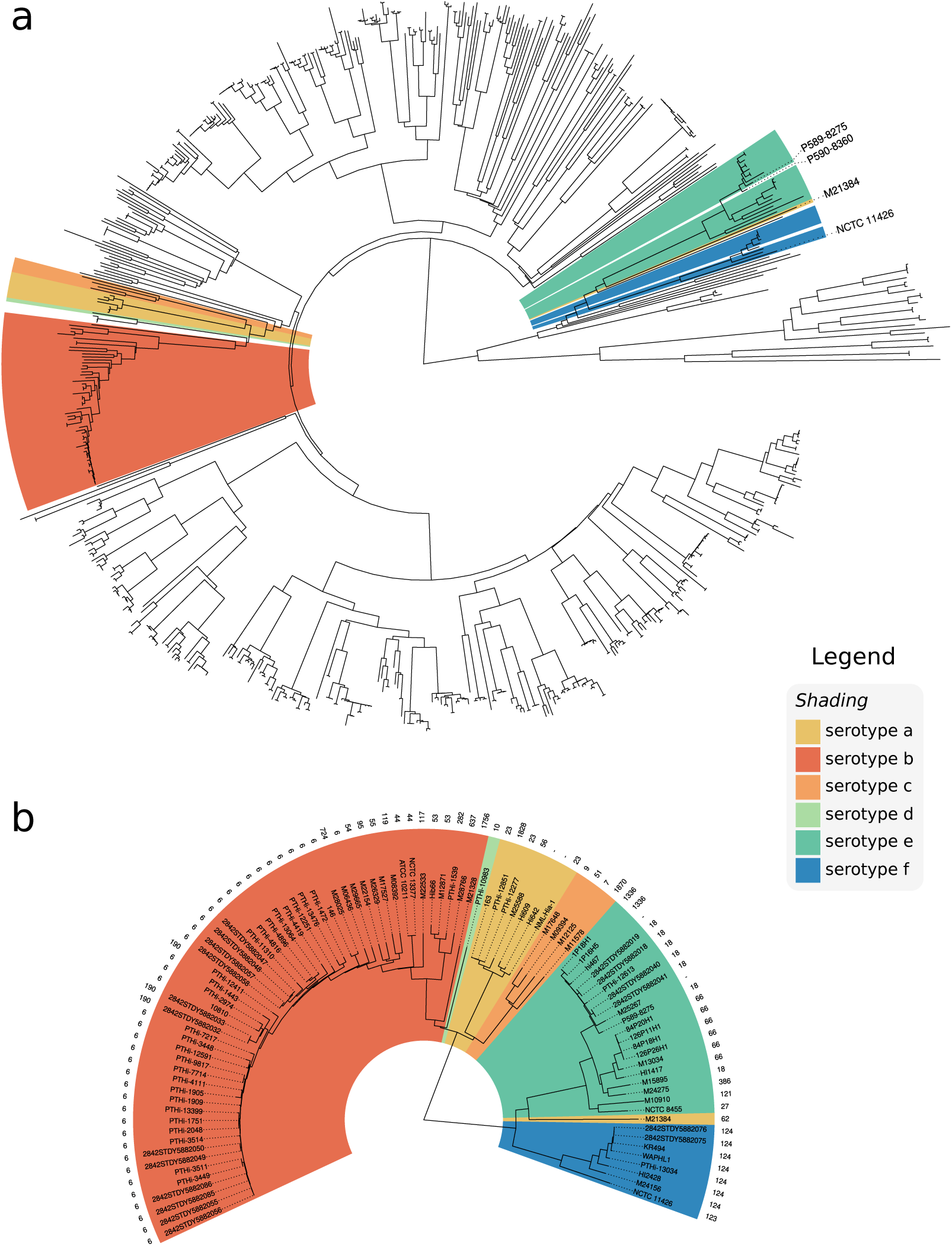
Whole genome neighbour-joining phylogeny inferred from MASH distances of assemblies in the GenBank dataset. Isolates are annotated with respective serotype as predicted by hicap. (a) Distribution of capsular serotypes in complete dataset. (b) The phylogeny subtree including only isolates that contained a *cap* locus additionally annotated with sequence type.

The whole genome phylogeny confirmed that encapsulated strains are relatively clonal and are generally restricted to serotype-specific monophyletic clades (Figure 4a), suggesting that each serotype emerged once within the *H. influenzae* population. This is consistent with earlier studies based on electrophoretic typing [35, 36], 16S rRNA [37], MLST [26], and WGS studies [38]. Here the whole genome phylogeny resolves the monophyletic nature of each capsule locus with respect to phylogenetic lineage on a larger scale and in greater detail.

hicap did not detect *cap* loci in a small number of isolates within these serotype-specific clades, indicating occasional capsule loss. For example P590-8360 clustered with the Hie clade (Figure 4) but no *cap* locus was identified by hicap or manual inspection of the assembly data. The high nucleotide identity between P590-8360 and the *cap*-e positive strain P589-8275 along the rest of the genome (see tree, Figure 4a) suggests that loss of the *cap*-e locus in the P590-8360 genome is the mostly likely explanation. Indeed the loss of capacity to synthesise capsule has previously been observed to occur by partial or complete deletion of the *cap* locus [38] and the rate of spontaneous capsule loss is estimated to occur at a frequency of 0.1 to 0.3% [39]. Our data is consistent with deletion of the *cap* locus being a cause of this phenomenon. Interestingly one *cap*-a genome (M21384) falls within the *cap*-e serotype-specific clade, suggesting possible recombination in this strain (Figure 4; further evidence for this is discussed below).

The serotype-specific clades cluster into two superclades within the *H. influenzae* phylogeny: one containing *cap* loci encoding Hia, Hib, Hic, and Hid and the other containing Hie and Hif *cap* loci (Figure 4). Individual gene trees for region I (*bex*) and III (*hcs*) genes show the same two-clade structure (Figure 5) as the core genomes of their host strains. This observation is consistent with diversification of these *cap* locus regions *in situ* within their host chromosomes following introduction into two distinct *H. influenzae* superclade ancestors. Whilst there is a general lack of homology between region II genes (Figure 1), two of the three pairs that do show a measure of homology (*cap*- c/*cap*-f and *cap*-d/*cap*-e) span both superclades, hence the evolutionary history of region II (and thus the distinct capsular serotypes) remains cryptic.

**Fig 5.**
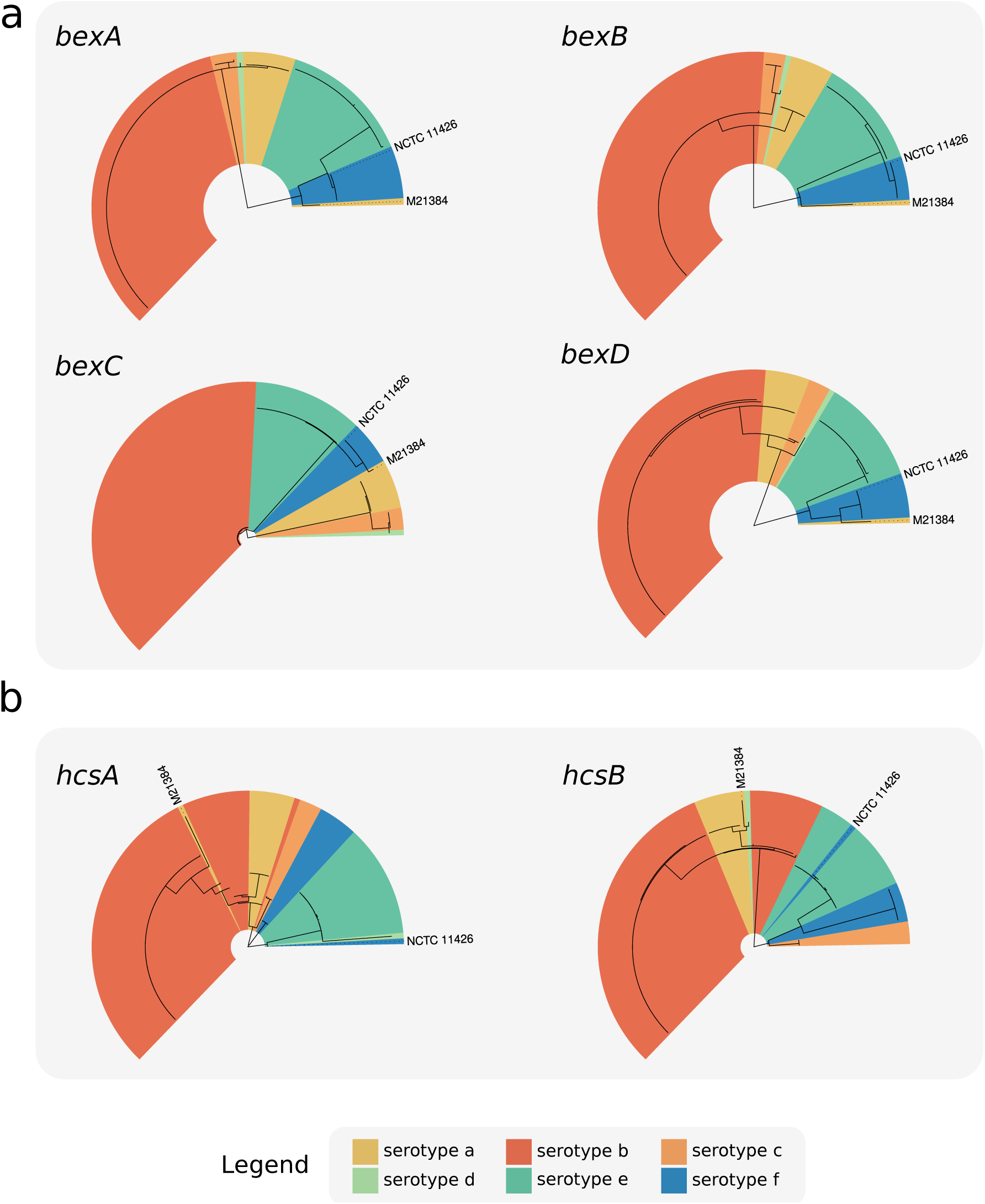
Phylogenies of all complete *cap* locus region (a) I and (b) III genes identified in the GenBank dataset. FastTree was used to recovery phylogenies from MAFFT gene nucleotide sequence alignments and isolates were annotated using serotype as predicted by hicap.

Variation in each region I or III gene was associated with serotype, suggesting that the sequence of any could potentially be used to predict capsule type with a relatively high degree of certainty (Figure 5). Indeed both *bexA* and *bexB* have been proposed and used in single gene PCR assays for the purpose of serotyping [40, 41]. Here the gene *bexB* showed the greatest differentiation between serotype-specific alleles (0.63% to 17.71% median pairwise nucleotide divergence; see Supplementary Figure S1) and contained only serotype-specific monophyletic clades in the gene tree. These data suggest *bexB* to be the most suitable single marker gene for use in PCR or sequenced-based prediction of serotype. In contrast, *bexA* showed less differentiation then *bexB*, particularly between serotypes a, b, c, and d (0.17% to 0.85% median pairwise nucleotide divergence).

The exceptions to the general association between region I/III genes and predicted serotype were two isolates, M21384 and NCTC 11426 (labelled in Figure 5). hicap predicted isolates M21384 and NCTC 11426 to be of serotype a and serotype f respectively. However both carry *cap* region I and/or III gene sequences distinct from other strains sharing the same *cap* II region type (and thus same predicted serotype), indicative of recombination involving *cap* locus within these isolates. Thus there is evidence for occasional recombination within the *cap* locus between the different serotype-specific variants, which would limit the accuracy of any single marker gene-based approach to serotype prediction.

### Recombination affecting the *cap* locus

The isolate M21384 was the only exception to clonal clustering by serotype in the whole genome phylogeny (Figure 4). While this isolate is predicted as Hia based on the presence of *cap*-a region II genes, the genome falls outside of the Hia clade and within the Hie/Hif superclade (Figure 4b). In all gene trees M21384 also did not cluster in the expected Hia serotype clade, suggesting there has been recombination within the *cap* locus of this isolate (see Figure 5). Similarly in the *hcsA* and *hcsB* gene trees, the isolate NCTC 11426 did not cluster with expect serotype Hif clade (Figure 5b). Given the phylogenetic relation of M21384 and NCTC 11426 to other capsular serotypes, it was suspected that these two strains result from recombination events affecting the *cap* locus (representing a 2% recombination rate involving the *cap* locus).

To better understand the recombinant *cap* loci in isolates M21384 and NCTC 11426 we first examined their positions in the whole genome phylogeny (Figure 4) and the *cap* locus gene trees (Figure 5), and then compared the full-length *cap* locus sequences of both isolates to reference *cap* locus sequences (Figure 6). NCTC 11426 (predicted Hif) belongs to the Hif clade in the whole genome tree and carries typical *cap*-f regions I and II but contains region III genes more similar to those from *cap*-e (Figure 6a). Hence it appears the *cap* locus of NCTC 11426 has resulted from a small recombination event between a Hif clade strain and the *cap* locus from a Hie clade strain. M21384 clusters within the Hie/Hif superclade of the whole genome phylogeny and carries *cap*-f-like region I genes (Figure 5a and 6b). However this isolate encodes *cap*-a like region II genes with a *cap*-b-like region III gene (*hcsA*) (Figure 5b and 6b). The gene content of M21384 *cap* locus suggests at least one recombination event involving import of foreign *cap* locus DNA into a Hif strain. It would be interesting to ascertain whether the isolates with recombinant *cap* loci described here do in fact express capsule and if so, to then establish serological phenotype. However serotyping has not been performed to our knowledge for either strain or they are not publicly available.

**Fig 6.**
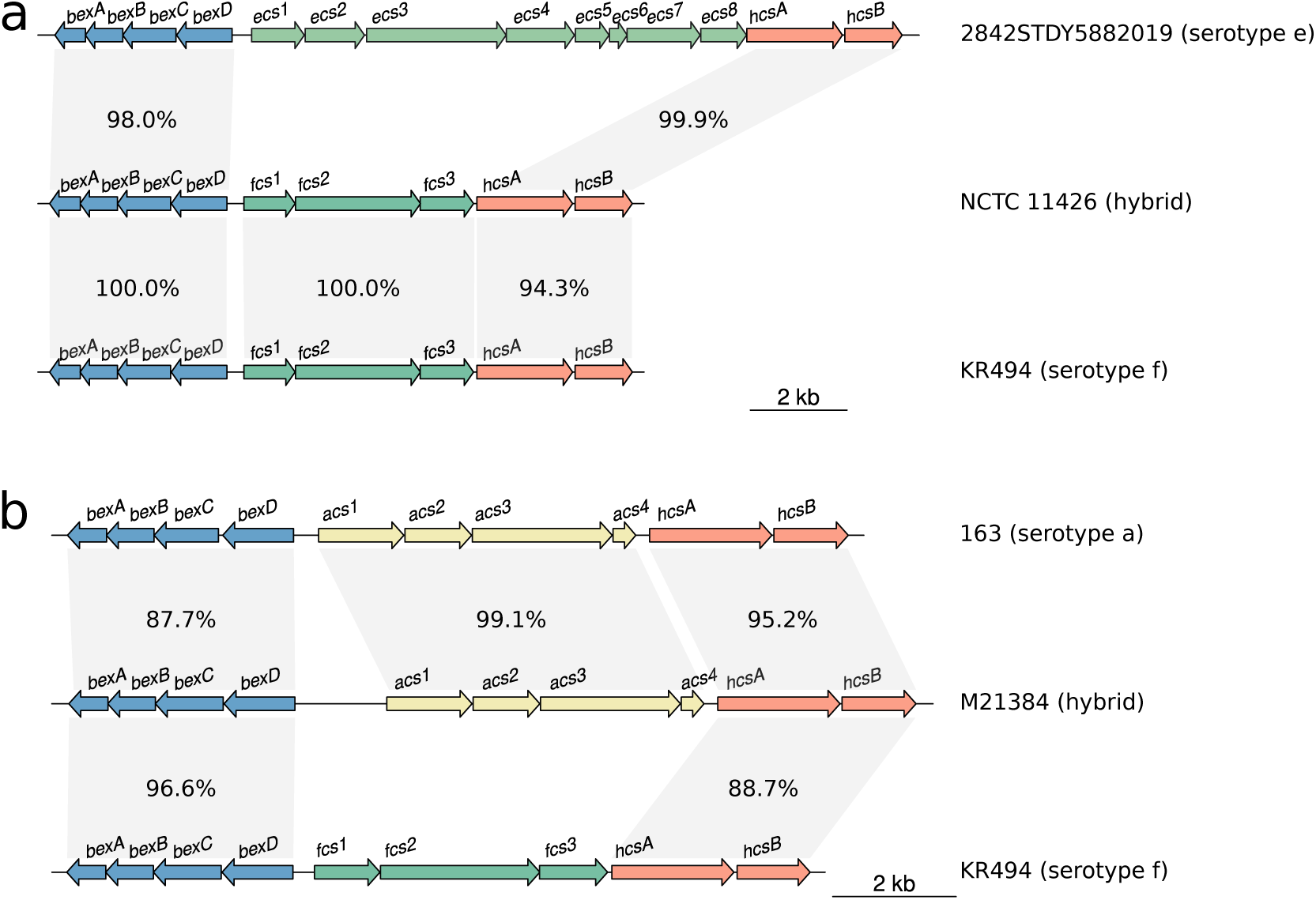
Homology plots generated using R and genoPlotR showing the *cap* locus of two isolates, which appear to have been subject to recombination. Different regions of (a) NCTC 11426 and (b) M21384 *cap* locus show varying homology to different reference *cap* loci suggesting a recombinogenic ancestry.

## Conclusion

The need for new tools and methods that leverage WGS continues to become increasingly pivotal with the adoption of WGS by public health laboratories. In this study we validated and demonstrated the robustness of hicap for prediction of *H. influenzae* serotype and capsule locus structure. The application of hicap to WGS enables rapid and accurate acquisition of capsule information to aid genomic studies at both an individual and population scale. We were also able to explore the diversity and distribution of *cap* loci in the *H. influenzae* population at unprecedented nucleotide resolution, identifying a likely misreported serotype in NCTC and describing two novel *H. influenzae cap* locus recombinants. The resurgence of disease caused by encapsulated *H. influenzae* and the potential for further antigenic diversification through recombination presents a potential public health issue. An important question is whether the geographically disparate reports of increasing cases of infection with non-Hib encapsulated strains reflects the emergence and wide dissemination of a small number of high successful disease-causing subclones (i.e. rare but worrying event) or multiple independent events reflecting sporadic but localised outbreaks of non-Hib disease. hicap will facilitate extracting answers to these and other questions from genomic surveillance data.

## Acknowledgments

This work was supported by the Bill & Melinda Gates Foundation, Seattle and Australian Government Research Training Program. KEH is supported by a Senior Medical Research Fellowship from the Viertel Foundation of Victoria.

All the authors declare that there are no conflicts of interest.

## Supporting information

S1 File. *H. influenzae* strains downloaded from GenBank (as of October 8, 2018). Data provided includes strain names, GenBank assembly accessions, predicted serotype, and other *cap* locus attributes.

---

**Fig S1.**
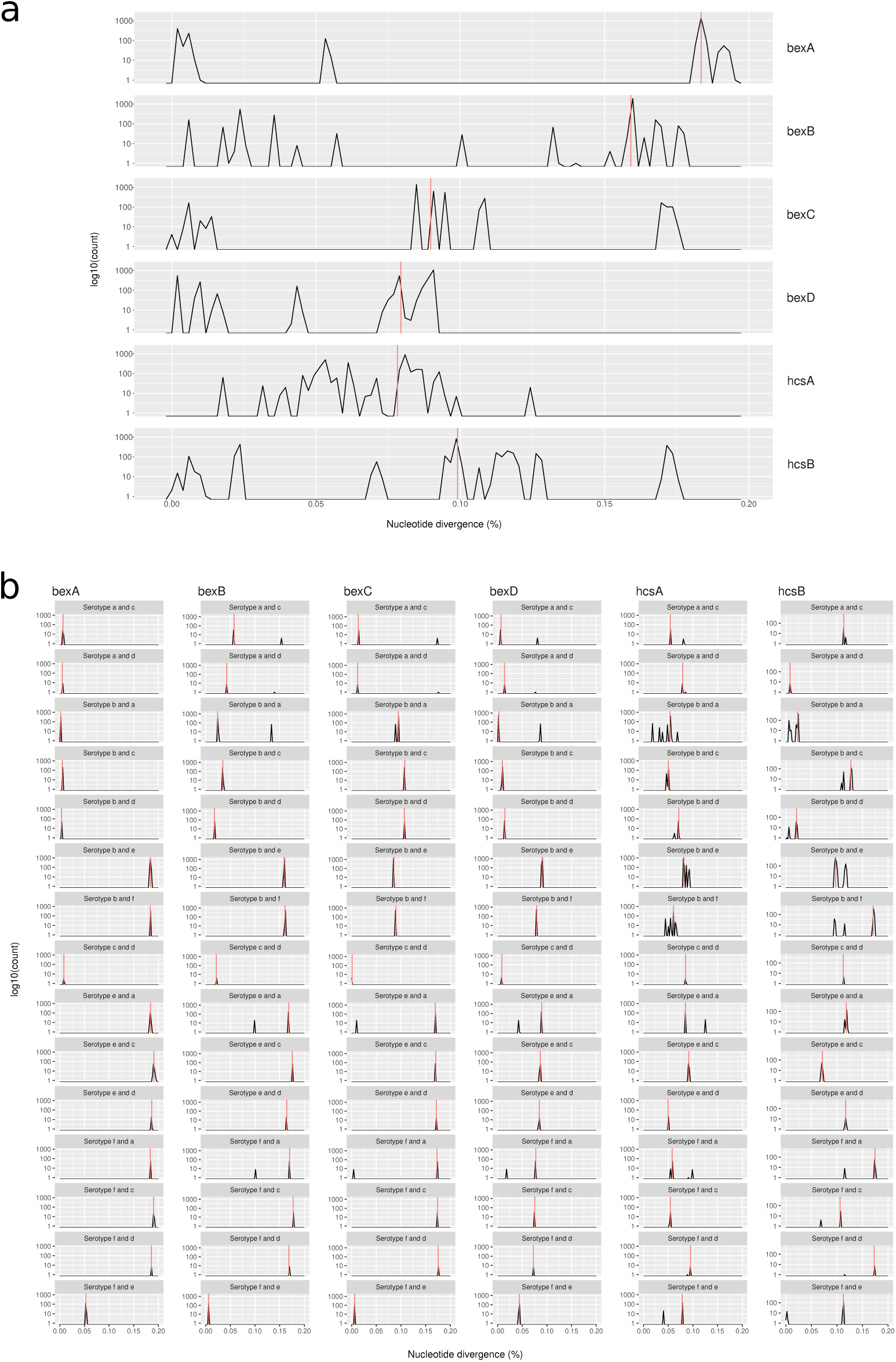
Distributions of nucleotide divergence measured for every gene sequence in different serotype groups from the GenBank dataset with median divergence indicated by red lines. Nucleotide divergence calculated by ape::dist.dna from MAFFT alignments for all *cap* region I and III gene sequences. (a) Combined distributions for all pair-wise nucleotide divergence measures of each gene. (b) Distribution of pair-wise measurements for each sequence between serotype groups of each gene.

## References

1. R. D. Fleischmann, M. D. Adams, O. White, R. A. Clayton, E. F. Kirkness, A. R. Kerlavage, C. J. Bult, J. F. Tomb, B. A. Dougherty, and J. M. Merrick, “Whole-genome random sequencing and assembly of *Haemophilus influenzae* Rd,” Science, vol. 269, no. 5223, pp. 496–512, 1995.

2. M. Pittman, “Variation and type specificity in the bacterial species *Hemophilus influenzae*,” The Journal of Experimental Medicine, vol. 53, no. 4, pp. 471–492, 1931.

3. J. S. Kroll, B. Loynds, L. N. Brophy, and E. R. Moxon, “The *bex* locus in encapsulated *Haemophilus influenzae*: a chromosomal region involved in capsule polysaccharide export,” Molecular Microbiology, vol. 4, no. 11, pp. 1853–1862, 1990.

4. S. Sukupolvi-Petty, S. Grass, and J. W. St Geme, “The *Haemophilus influenzae* Type b *hcsA* and *hcsB* gene products facilitate transport of capsular polysaccharide across the outer membrane and are essential for virulence,” Journal of Bacteriology, vol. 188, no. 11, pp. 3870–3877, 2006.

5. A. Follens, M. Veiga-da Cunha, R. Merckx, E. van Schaftingen, and J. van Eldere, “*acs1* of *Haemophilus influenzae* type a capsulation locus region II encodes a bifunctional ribulose 5-phosphate reductase- CDP-ribitol pyrophosphorylase,” Journal of Bacteriology, vol. 181, no. 7, pp. 2001–2007, 1999.

6. J. Van Eldere, L. Brophy, B. Loynds, P. Celis, I. Hancock, S. Carman, J. S. Kroll, and E. R. Moxon, “Region II of the *Haemophilus influenzae* type be capsulation locus is involved in serotype-specific polysaccharide synthesis,” Molecular Microbiology, vol. 15, no. 1, pp. 107–118, 1995.

7. T.-T. Lâm, H. Claus, M. Frosch, and U. Vogel, “Sequence analysis of serotype-specific synthesis regions II of *Haemophilus influenzae* serotypes c and d: evidence for common ancestry of capsule synthesis in *Pasteurellaceae* and *Neisseria meningitidis*,” Research in Microbiology, vol. 162, no. 5, pp. 483–487, 2011.

8. M. Giufrè., R. Cardines, P. Mastrantonio, and M. Cerquetti, “Genetic Characterization of the Capsulation Locus of *Haemophilus influenzae* Serotype e,” Journal of Clinical Microbiology, vol. 48, no. 4, pp. 1404–1407, 2010.

9. S. W. Satola, P. L. Schirmer, and M. M. Farley, “Genetic Analysis of the Capsule Locus of *Haemophilus influenzae* Serotype f,” Infection and Immunity, vol. 71, no. 12, pp. 7202–7207, 2003.

10. P. G. Corn, J. Anders, A. K. Takala, H. Käyhty, and S. K. Hoiseth, “Genes involved in *Haemophilus influenzae* type b capsule expression are frequently amplified,” The Journal of Infectious Diseases, vol. 167, no. 2, pp. 356–364, 1993.

11. J. S. Kroll, I. Hopkins, and E. R. Moxon, “Capsule loss in *H. influenzae* type b occurs by recombination-mediated disruption of a gene essential for polysaccharide export,” Cell, vol. 53, no. 3, pp. 347–356, 1988.

12. B. G. Kapogiannis, S. Satola, H. L. Keyserling, and M. M. Farley, “Invasive infections with *Haemophilus influenzae* serotype a containing an IS*1016*-*bexA* partial deletion: possible association with virulence,” Clinical Infectious Diseases: An Official Publication of the Infectious Diseases Society of America, vol. 41, no. 11, pp. e97–103, 2005.

13. H. A. Bijlmer, “World-wide epidemiology of *Haemophilus influenzae* meningitis; industrialized versus non-industrialized countries,” Vaccine, vol. 9 Suppl, pp. S5–9; discussion S25, 1991.

14. H. Peltola, “Worldwide *Haemophilus influenzae* type b disease at the beginning of the 21st century: global analysis of the disease burden 25 years after the use of the polysaccharide vaccine and a decade after the advent of conjugates,” Clinical Microbiology Reviews, vol. 13, no. 2, pp. 302–317, 2000.

15. M. Ulanova, “Global Epidemiology of Invasive *Haemophilus influenzae* Type a Disease: Do We Need a New Vaccine?,” Journal of Vaccines, vol. 2013, p. 14, 2013.

16. V. Eton, A. Schroeter, L. Kelly, M. Kirlew, R. S. W. Tsang, and M. Ulanova, “Epidemiology of invasive pneumococcal and *Haemophilus influenzae* diseases in Northwestern Ontario, Canada, 2010-2015,” International journal of infectious diseases: IJID: official publication of the International Society for Infectious Diseases, vol. 65, pp. 27–33, 2017.

17. C. Puig, I. Grau, S. Marti, F. Tubau, L. Calatayud, R. Pallares, J. Liñares, and C. Ardanuy, “Clinical and Molecular Epidemiology of *Haemophilus influenzae* Causing Invasive Disease in Adult Patients,” PLOS ONE, vol. 9, no. 11, p. e112711, 2014.

18. J. Revez, L. Espinosa, B. Albiger, K. C. Leitmeyer, M. J. Struelens, and ECDC National Microbiology Focal Points and Experts Group, “Survey on the Use of Whole-Genome Sequencing for Infectious Diseases Surveillance: Rapid Expansion of European National Capacities, 2015-2016,” Frontiers in Public Health, vol. 5, p. 347, 2017.

19. J. Besser, H. A. Carleton, P. Gerner-Smidt, R. L. Lindsey, and E. Trees, “Next-generation sequencing technologies and their application to the study and control of bacterial infections,” Clinical Microbiology and Infection: The Official Publication of the European Society of Clinical Microbiology and Infectious Diseases, vol. 24, no. 4, pp. 335–341, 2018.

20. Y. H. Grad and M. Lipsitch, “Epidemiologic data and pathogen genome sequences: a powerful synergy for public health,” Genome Biology, vol. 15, no. 11, p. 538, 2014.

21. D. Hyatt, G.-L. Chen, P. F. Locascio, M. L. Land, F. W. Larimer, and L. J. Hauser, “Prodigal: prokaryotic gene recognition and translation initiation site identification,” BMC bioinformatics, vol. 11, p. 119, 2010.

22. C. Camacho, G. Coulouris, V. Avagyan, N. Ma, J. Papadopoulos, K. Bealer, and T. L. Madden, “BLAST+: architecture and applications,” BMC Bioinformatics, vol. 10, p. 421, 2009.

23. P. J. A. Cock, T. Antao, J. T. Chang, B. A. Chapman, C. J. Cox, A. Dalke, I. Friedberg, T. Hamelryck, F. Kauff, B. Wilczynski, and M. J. L. de Hoon, “Biopython: freely available Python tools for computational molecular biology and bioinformatics,” Bioinformatics, vol. 25, no. 11, pp. 1422–1423, 2009.

24. A. Bankevich, S. Nurk, D. Antipov, A. A. Gurevich, M. Dvorkin, A. S. Kulikov, V. M. Lesin, S. I. Nikolenko, S. Pham, A. D. Prjibelski, A. V. Pyshkin, A. V. Sirotkin, N. Vyahhi, G. Tesler, M. A. Alekseyev, and P. A. Pevzner, “SPAdes: A New Genome Assembly Algorithm and Its Applications to Single-Cell Sequencing,” Journal of Computational Biology, vol. 19, no. 5, pp. 455–477, 2012.

25. B. D. Ondov, T. J. Treangen, P. Melsted, A. B. Mallonee, N. H. Bergman, S. Koren, and A. M. Phillippy, “Mash: fast genome and metagenome distance estimation using MinHash,” Genome Biology, vol. 17, no. 1, p. 132, 2016.

26. E. Meats, E. J. Feil, S. Stringer, A. J. Cody, R. Goldstein, J. S. Kroll, T. Popovic, and B. G. Spratt, “Characterization of encapsulated and noncapsulated *Haemophilus influenzae* and determination of phylogenetic relationships by multilocus sequence typing,” Journal of Clinical Microbiology, vol. 41, no. 4, pp. 1623–1636, 2003.

27. L. Guy, J. Roat Kultima, and S. G. E. Andersson, “genoPlotR: comparative gene and genome visualization in R,” Bioinformatics, vol. 26, no. 18, pp. 2334–2335, 2010.

28. R. C. Team, *R: A Language and Environment for Statistical Computing*. Vienna, Austria: R Foundation for Statistical Computing, 2019.

29. G. Yu, D. K. Smith, H. Zhu, Y. Guan, and T. T.-Y. Lam, “ggtree: an R package for visualization and annotation of phylogenetic trees with their covariates and other associated data,” Methods in Ecology and Evolution, vol. 8, no. 1, pp. 28–36, 2017.

30. K. Katoh and D. M. Standley, “MAFFT multiple sequence alignment software version 7: improvements in performance and usability,” Molecular Biology and Evolution, vol. 30, no. 4, pp. 772–780, 2013.

31. M. N. Price, P. S. Dehal, and A. P. Arkin, “FastTree 2 – Approximately Maximum-Likelihood Trees for Large Alignments,” PLOS ONE, vol. 5, no. 3, p. e9490, 2010.

32. E. Paradis, J. Claude, and K. Strimmer, “APE: Analyses of Phylogenetics and Evolution in R language,” Bioinformatics, vol. 20, no. 2, pp. 289–290, 2004.

33. S. W. Satola, J. T. Collins, R. Napier, and M. M. Farley, “Capsule gene analysis of invasive *Haemophilus influenzae*: accuracy of serotyping and prevalence of IS*1016* among nontypeable isolates,” Journal of Clinical Microbiology, vol. 45, no. 10, pp. 3230–3238, 2007.

34. J. S. Kroll, E. R. Moxon, and B. M. Loynds, “An ancestral mutation enhancing the fitness and increasing the virulence of *Haemophilus influenzae* type b,” The Journal of Infectious Diseases, vol. 168, no. 1, pp. 172–176, 1993.

35. J. M. Musser, D. M. Granoff, P. E. Pattison, and R. K. Selander, “A population genetic framework for the study of invasive diseases caused by serotype b strains of *Haemophilus influenzae*,” Proceedings of the National Academy of Sciences of the United States of America, vol. 82, no. 15, pp. 5078–5082, 1985.

36. J. M. Musser, J. S. Kroll, D. M. Granoff, E. R. Moxon, B. R. Brodeur, J. Campos, H. Dabernat, W. Frederiksen, J. Hamel, and G. Hammond, “Global genetic structure and molecular epidemiology of encapsulated *Haemophilus influenzae*,” Reviews of Infectious Diseases, vol. 12, no. 1, pp. 75–111, 1990.

37. C. T. Sacchi, D. Alber, P. Dull, E. A. Mothershed, A. M. Whitney, G. A. Barnett, T. Popovic, and L. W. Mayer, “High level of sequence diversity in the 16S rRNA genes of *Haemophilus influenzae* isolates is useful for molecular subtyping,” Journal of Clinical Microbiology, vol. 43, no. 8, pp. 3734–3742, 2005.

38. M. De Chiara, D. Hood, A. Muzzi, D. J. Pickard, T. Perkins, M. Pizza, G. Dougan, R. Rappuoli, E. R. Moxon, M. Soriani, and C. Donati, “Genome sequencing of disease and carriage isolates of nontypeable *Haemophilus influenzae* identifies discrete population structure,” Proceedings of the National Academy of Sciences of the United States of America, vol. 111, no. 14, pp. 5439–5444, 2014.

39. S. K. Hoiseth, C. J. Connelly, and E. R. Moxon, “Genetics of spontaneous, high-frequency loss of b capsule expression in *Haemophilus influenzae*,” Infection and Immunity, vol. 49, no. 2, pp. 389–395, 1985.

40. G. S. Davis, S. A. Sandstedt, M. Patel, C. F. Marrs, and J. R. Gilsdorf, “Use of *bexB* to detect the capsule locus in *Haemophilus influenzae*,” Journal of Clinical Microbiology, vol. 49, no. 7, pp. 2594–2601, 2011.

41. T. J. Falla, D. W. Crook, L. N. Brophy, D. Maskell, J. S. Kroll, and E. R. Moxon, “PCR for capsular typing of *Haemophilus influenzae*,” Journal of Clinical Microbiology, vol. 32, no. 10, pp. 2382–2386, 1994.

42. Y.-C. Su, F. Hörhold, B. Singh, and K. Riesbeck, “Complete Genome Sequence of Encapsulated *Haemophilus influenzae* Type f KR494, an Invasive Isolate That Caused Necrotizing Myositis,” Genome Announcements, vol. 1, no. 5, pp. e00470–13, 2013.

43. M. Staples, R. M. A. Graham, and A. V. Jennison, “Characterisation of invasive clinical *Haemophilus influenzae* isolates in Queensland, Australia using whole-genome sequencing,” Epidemiology and Infection, vol. 145, no. 8, pp. 1727–1736, 2017.

44. M. Pittman, “Antibacterial Action of Several Sulfonamide Compounds on *Hemophilus influenzae* Type B,” Public Health Reports, vol. 57, no. 50, pp. 1899–1910, 1942.

45. M. Giufrè, R. Cardines, and M. Cerquetti, “First Whole-Genome Sequence of a *Haemophilus influenzae* Type e Strain Isolated from a Patient with Invasive Disease in Italy,” Genome Announcements, vol. 5, no. 13, 2017.

46. S. R. Dobson, J. S. Kroll, and E. R. Moxon, “Insertion sequence IS*1016* and absence of *Haemophilus* capsulation genes in the Brazilian purpuric fever clone of *Haemophilus influenzae* biogroup *aegyptius*,” Infection and Immunity, vol. 60, no. 2, pp. 618–622, 1992.

